# Contribution of DNA breathing to physical interactions with transcription factors

**DOI:** 10.1101/2025.01.20.633840

**Authors:** Waqaas Butt, Ben Lai, Tsu-Pei Chiu, Manish Bhattarai, Sheng Qian, Alan R. Bishop, Jubao Duan, Boian S. Alexandrov, Remo Rohs, Xin He

## Abstract

Interaction between transcription factors (TFs) and DNA plays a key role in regulating gene expression. It is generally believed that these interactions are controlled through recognition of DNA core motifs by TFs. Nevertheless, several studies pointed out the limitation of this view, in particular, DNA sequence variants influencing TF binding are often located outside of core motifs. One possible explanation is that the physical properties of DNA may play a role in TF-DNA interactions. Recent studies have supported the importance of DNA shape features, especially in flanking regions of core motifs. Another important physical property of DNA is DNA breathing, the spontaneous opening of double-stranded DNA through thermal motions. But there have been few genomic studies of the role of DNA breathing in TF-DNA interactions. In this work, we analyzed in vitro TF-DNA binding data of three TFs and found that DNA breathing features inside or near core motifs are correlated with binding affinity. This suggests that these TFs may prefer locally and temporally melted DNA formed through breathing. We extended the analysis to 44 TFs with in vivo ChIP-seq binding data. We found that for a large proportion of TFs, their breathing features in or near core motifs are associated with binding, but the sign and magnitude of these associations vary substantially across TF families. Altogether, our study supports the hypothesis that DNA breathing features near binding motifs contribute to TF-DNA interactions.

**Author Summary:** Proper regulation of when and where genes are expressed is crucial to biological development and function. This process is largely controlled by interaction of transcription factors (TFs) with DNA sequences. The recognition of specific DNA sequences by TFs is important to ensure that only the correct genes are activated. Extensive work has shown that TFs prefer to bind certain DNA sequence patterns of 6-20 bp, known as motifs. However, the structure of DNA molecules may also play a role. In this work, we explored the role of DNA breathing, which refers to spontaneous opening of double strand DNA due to thermal motions. This process creates transient, single-strand “bubbles” in DNA. Through examining TF-DNA binding data of >60 TFs, we found that the propensity of DNA forming bubbles near motifs is often associated with binding affinity of DNA sequence. Interestingly the patterns of these associations seem to vary with TFs. Altogether, our results highlighted the potential of DNA breathing in influencing TF-DNA interactions.

## Introduction

Spatial-temporal expression of genes is critical for proper development and functions of organisms. Gene expression is regulated, to a large extent, by the interactions of transcription factors (TFs) with their DNA target sites. A central problem in the field of gene regulation is thus to understand how these physical interactions form. It is widely believed that TFs recognize sequence-specific features, known as core motifs. Various methods have been developed to predict binding of TFs to putative regulatory sequences using motif features [1]. While the earlier methods focus on modeling interaction of TFs with a single binding site, recent methods, using neural networks, have attempted to capture complex, non-linear patterns of motifs within DNA sequences [2, 3, 4]. Despite the progress, there is still an important gap in our understanding of how TFs interact with DNA. From genetic studies, it was found that DNA variants that affect regulatory functions often reside outside known motifs [5, 6] One study showed, for example, that while 60% of putative causal variants of autoimmune diseases are located in enhancers, only 10-20% directly alter TF motifs [7]. Another study mapped DNA variants associated with TF binding, and found that only in 1% of cases, these DNA variants are located in predicted motifs. These studies thus strongly suggest that features of DNA sequences other than motifs contribute to interactions with TFs.

To fill this gap, researchers have proposed that DNA variants may alter epigenomic features of DNA sequences, thus their regulatory functions [8, 9]. For example, DNA variants may change DNA methylation, which can influence TF binding [10]. Other possibilities include GC content near TF binding sites and nucleosome positioning. Nevertheless, there is limited evidence that directly shows that alternation of these features by DNA variants leads to a qualitative change of TF binding. An alternative explanation lies in the physics of DNA itself. Indeed, in an important line of work, researchers found that DNA shape features near motifs play an important role in determining TF-DNA interactions. One study showed that incorporating DNA shape features at and near motifs improved the prediction of TF binding from in vitro studies [11]. Another study hypothesized a concept of DNA shape motifs, analogous to sequence motifs but based entirely on DNA shape features. These shape motifs may occur outside sequence motifs and were estimated to explain 17% of TF binding in ChIP-seq data [12]. More generally, these results together support the importance of DNA physical properties in determining TF binding.

DNA shape is not the only factor in determining TF binding, because TFs require hydrogen bonds for motif recognition [13]. Other physical characteristics of DNA double helices are probably important. One important feature is DNA breathing. This refers to spontaneous and transient opening of intra-base pair hydrogen bonds that form double-stranded DNA through thermal motions. This transient opening may lead to so-called DNA “bubbles” (Fig. 1a, right panel). DNA breathing dynamics is generally sequence-specific, with the probability of DNA bubble formation and other breathing features dependent on local sequence contexts. One reason for this is the composition of A/T versus G/C base pairs, which determines the strength of hydrogen bonding between the two DNA strands. Earlier studies showed that DNA breathing dynamics can influence TF binding [14, 15]. Importantly, sequence variants influencing DNA breathing may affect TF binding and regulatory functions of DNA. These earlier studies, however, were limited to a small number of TFs or DNA variants. A recent paper showed that incorporating DNA breathing features in sequence-only models provides a modest improvement of predicting TF binding targets [16]. However, this study has several limitations. Given the complex and non-linear nature of deep neural networks, the models from the study were difficult to interpret, and it remains unclear how DNA breathing may contribute to binding. Additionally, the performance gain seems to depend on the architecture of the neural networks: the best-performing sequence-only model has comparable performance with the model incorporating both sequence and DNA breathing.

**Fig. 1:**
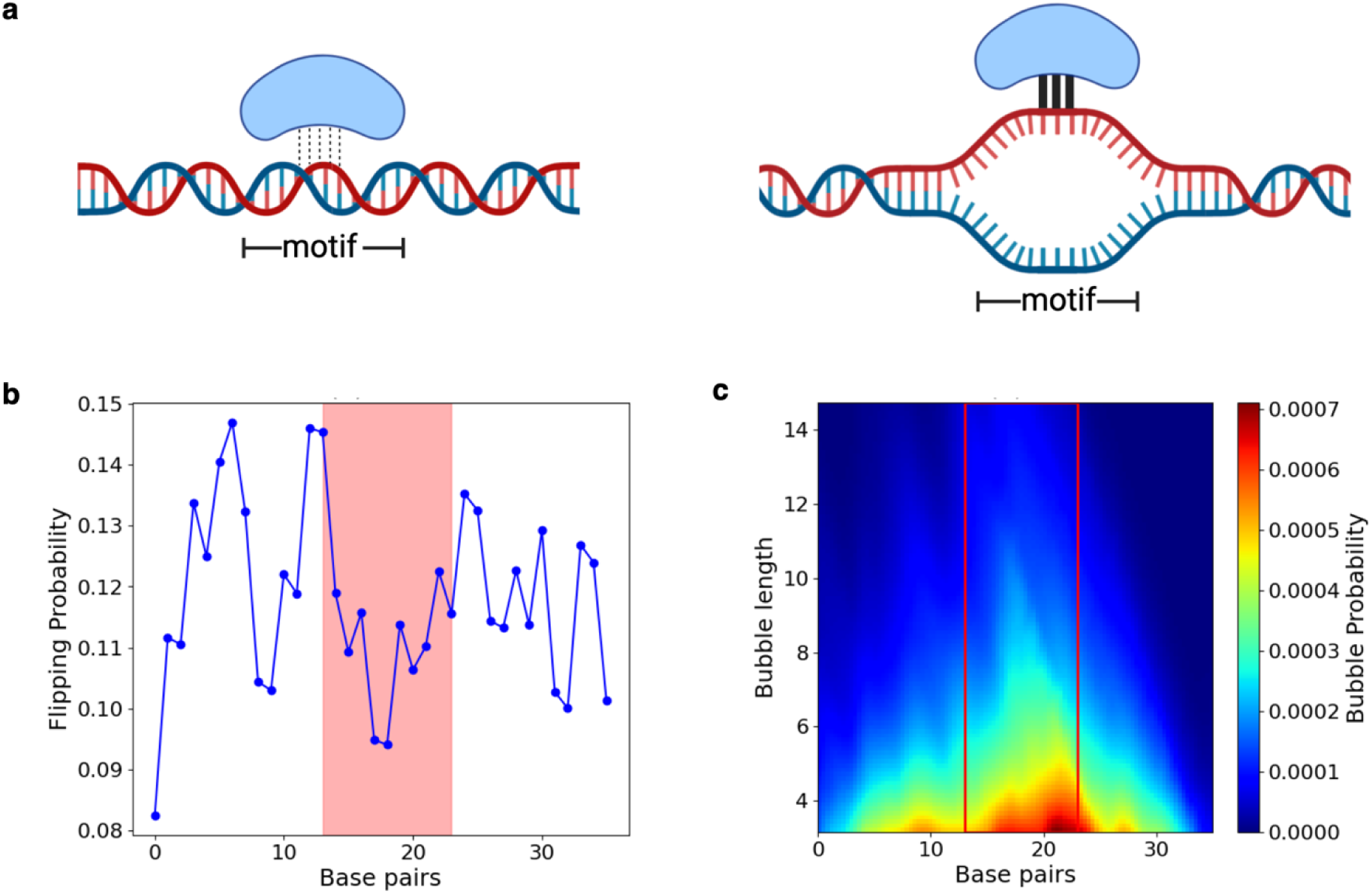
DNA breathing feature representation and TF-DNA binding hypothesis. (a) Our hypothesis states that TF binding to a motif can be affected by the presence of a bubble. (b) Base-flipping features for a single sequence calculated at a coordinate displacement threshold of about 0.5 * √2 Å. The red box indicates nucleotide positions where a TF core motif is located. (c) A heatmap of the bubble propensity feature generated for the same gcPBM sequence at a 3.5 Å amplitude threshold.

Our goal in this study was to perform a comprehensive analysis, based on statistical patterns of DNA breathing features and TF motif features, to examine the role of DNA breathing on TF binding. Our main hypothesis is that DNA breathing features, inside or near TF motifs, may influence the strength of TF-DNA interactions (Fig. 1a). For example, a DNA bubble, by opening adjacent DNA base pairs transiently, may facilitate binding of a TF. We think this is a plausible model. While TFs bind double-stranded DNA mostly in the major or minor groove, the presence of a bubble opens the base pairs of the core motif and allows additional or alternate interactions with the base-pairing edge of the nucleotides and potentially amino acid-base stacking, potentially improving binding affinity. We used two categories of DNA breathing features derived from DNA breathing dynamics based on physical simulations [17]. The first feature captures the local base-pair level breathing profile, while the second feature captures the breathing profile over an extended sequence window. Specifically, because of DNA breathing, a base pair may be transiently flipped outside of the double helix, and the probability of this event happening is denoted as ‘flipping probability’ feature, our first feature. (Fig. 1b, S1 Fig. e). At the extended sequence level, DNA breathing may create a bubble that spans multiple base-pair positions. Our second feature, “bubble propensity”, captures the probability of DNA bubble formation at any location of a given nucleotide sequence, thereby describing bubble location and length (Fig. 1c, S1 Fig. a). We note that bubble formation probability depends on the length and the width threshold we used in defining bubbles (see Methods). Both features capture the tendency of opening intra-base pair hydrogen bonds leading to melting of the double helix.

In our study, we used both features to examine the role of DNA breathing in TF-DNA interactions. We utilized two datasets to test our hypothesis. Genomic-context protein-binding microarray (gcPBM) is an in vitro technique to measure the strength of TF interaction with a large number of short DNA fragments of about 35 base pairs in length [18]. The advantage of using in vitro data is that it is solely based on physical interactions, without the complexity introduced by epigenomic and cellular features, e.g., chromatin accessibility [19] or TF concentration in the cell [20], that can influence TF binding. Compared to universal PBM (uPBM) which measures TF binding to tens of thousands of DNA oligomers, gcPBM arrays are longer and, more importantly, contain the flanking sequences of tested DNA oligomers form actual genomic context. The gcPBM dataset, however, was limited to only three TFs from the basis helix-loop-helix (bHLH) family. We thus used a second, in vivo, dataset consisting of ChIP-seq data of many more TFs from different protein families.

## Results

### DNA flipping probability feature correlates with TF binding in gcPBM data

We first examined the relationship of DNA breathing and TF binding using the flipping probability feature. We hypothesized that flipping probability profiles at the motif locations would influence binding of TFs. To test this, we used gcPBM data of three bHLH TF complexes: *Max-Max homodimer (MAX)*, *Mad2-Max heterodimer (MAD)*, and *c-Myc-Max heterodimer (MYC)* [18] (S1 Table). We focused on the results of *MAX* and *MYC* complexes here, given that *MAD* did not show a similarly well-defined motif. For each TF complex, we considered all sequences that contain at least a match to the TF motif. For any such sequence, we computed the average flipping probability across all nucleotide positions in the motif match. When a sequence contains multiple motif matches, all the matched positions will be included for computing flipping probability.

We observed highly significant correlations between average flipping probability and DNA binding affinity for both *MAX* and *MYC* (Fig 2a). The correlations are positive in both cases, suggesting that sequences with higher average flipping probability at motif sites tended to exhibit higher binding affinity. Using other ways of defining flipping features, using the relative base displacement within base pairs, gave similar results (S2 Fig).

**Fig. 2:**
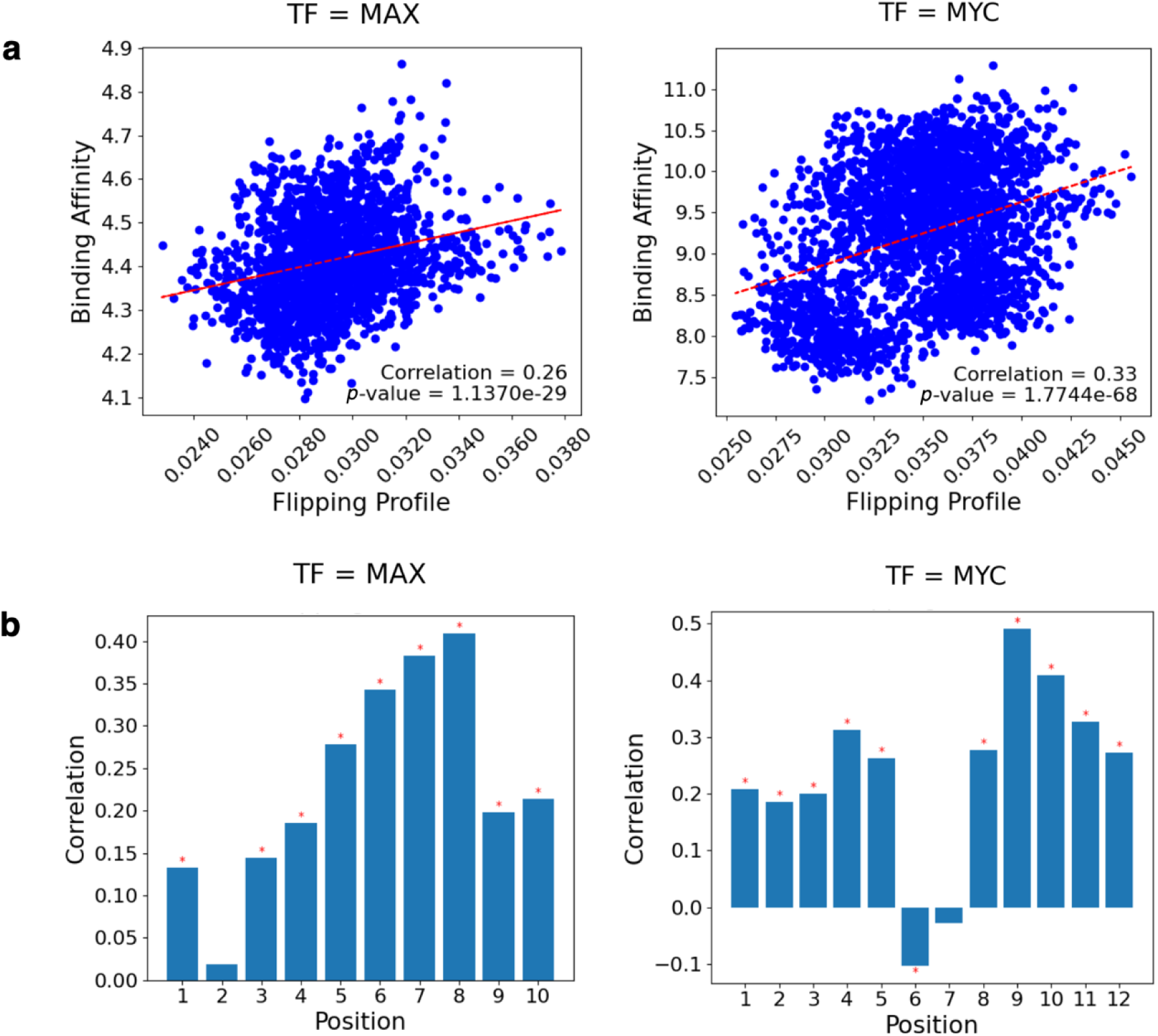
Correlations of DNA flipping features with binding affinity at motif locations. Nucleotide positions 3-8 for *MAX* and 4-9 for *MYC* represent the E-box motif of bHLH proteins. (a) The average flipping probability of nucleotide positions was correlated with binding affinity measured by gcPBM for *MAX* and *MYC*. (b) The flipping probability at each position in the motif for *MAX* and *MYC* were independently correlated with binding affinity. Bonferroni corrected correlation significance with *ɑ* = 0.05 is denoted by a red asterisk.

We next hypothesized that DNA flipping features at certain locations in a motif may be particularly important for determining TF-DNA interactions. We thus examined the correlation of flipping probability at each nucleotide position against binding affinity (Fig 2b). For both TFs, we observed substantial variations of the correlation across motif positions. For *MAX*, the correlation is not significant at position 2, and reached its peak at position 8, indicating that flipping at this position is likely influential on binding affinity. *MYC* also demonstrated a complex pattern, with a notable peak in correlation near position 9, while position 6 exhibited a slight negative correlation. Altogether, these results suggest that flipping profiles at most positions in a motif contribute positively to binding, while flipping features at a small number of positions have a nuanced role in the binding dynamics of *MAX* and *MYC*.

### DNA bubble propensity correlates with TF binding affinities in gcPBM data

As an alternative strategy of studying the role of DNA breathing in TF binding, we examined whether the presence of bubble features within DNA target regions correlates with binding affinity for *MAD*, *MAX*, and *MYC*. Across all three TFs, sequences with detectable bubbles (denoted as bubble positive sequences) generally exhibited higher binding affinity compared to those without detectable bubbles (bubble negative) (Fig 3a). The difference in the distributions of binding affinities were statistically significant for all three TFs, based on Wilcoxon rank sum test. These results suggest that the presence of DNA bubbles is associated with higher TF binding affinity, supporting our hypothesis that bubbles enhance binding.

**Fig. 3:**
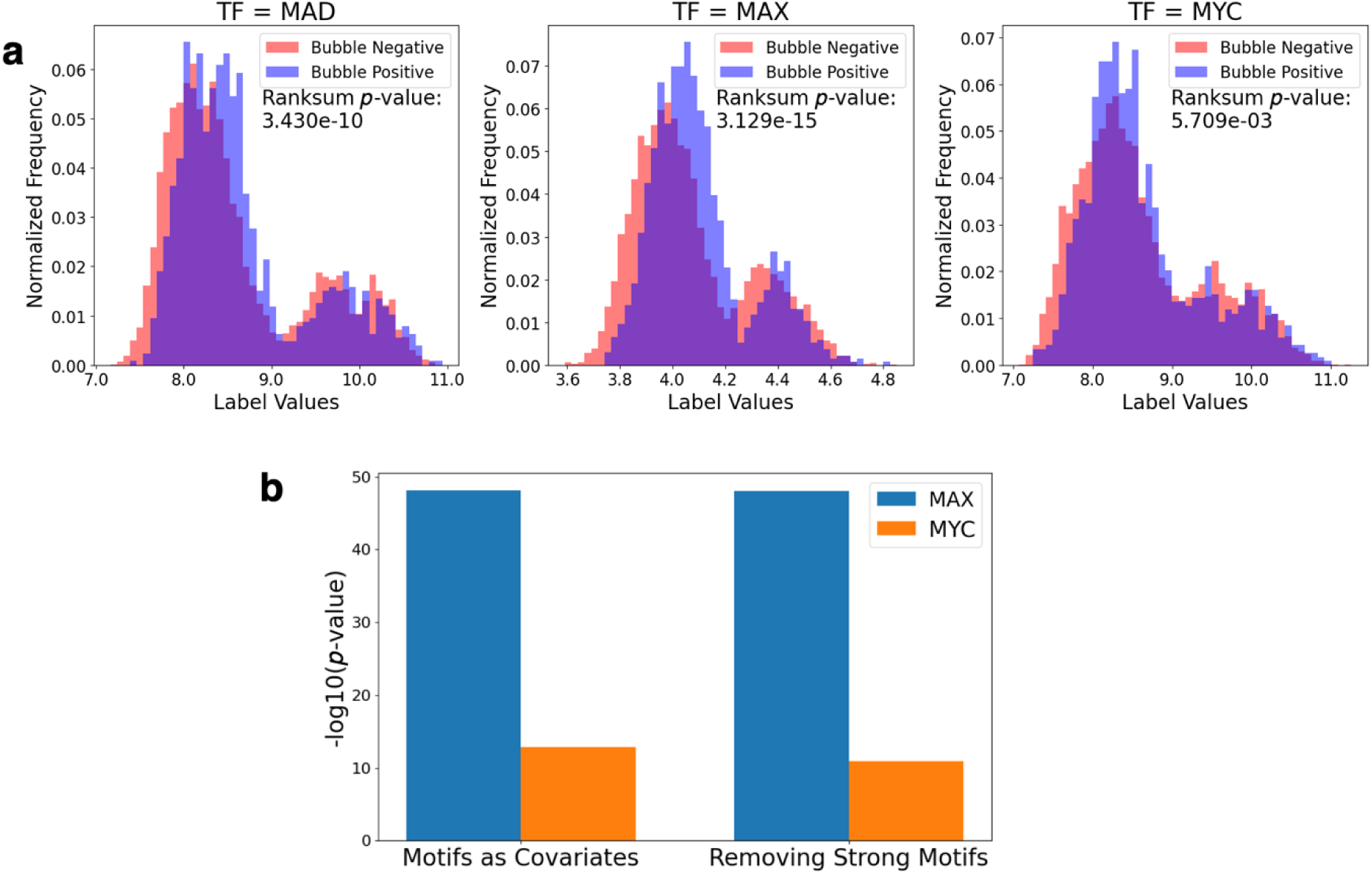
Binding affinity correlations with bubble presence. (a) Histogram of the binding affinity of TF-bound sequences in the bHLH gcPBM dataset (*MAD*, *MAX*, *MYC*) split according to bubble presence or absence, with normalized frequencies. Wilcoxon rank sum test performed to assess statistical significance of the difference in distributions of the two groups. (b) Linear regression significance values of bubble propensity features fitted against binding affinity with different strategies to control for the existence of motifs.

This analysis, however, is complicated by the presence of motifs in DNA target sequences. Indeed, the number and strength of motifs is a major factor determining TF-DNA interactions and may confound the relationship of DNA bubble propensity and binding affinity. To further explore the relationship between bubble features and binding affinity, we conducted linear regression analyses incorporating different strategies to control for the effect of motifs. We included motif presence as covariates in the regression model or removed sequences with strong motifs from the analysis. In each case, the bubble features remained significantly correlated with binding affinity for both *MAX* and *MYC* (Fig. 3b).

### DNA breathing features correlate with binding of some TFs in ChIP-seq data

To gain additional insight into how DNA breathing influences TF binding in complex cellular context, we extended our analysis to a much larger in vivo TF binding dataset derived from ChIP-seq. This dataset was collected from ChIP-seq studies of 64 TFs [2]. Among these TFs, 44 have known binding motifs, so our analysis focused on these 44 TFs (S2 Table). Our goal was to test if the patterns observed with the bHLH gcPBM dataset [18] can be replicated, and whether the role of DNA breathing in TF-DNA interaction varies with specific TF families. For the ChIP-seq data of each TF, we classified the peaks in the data as TF-bound sequences. We only included TFs in the analysis that have at least 3000 peaks. We also defined a large set of negative-control sequences where TFs were not bound to DNA (see Methods). We then performed a similar analysis as for the gcPBM data using the flipping probability feature: we correlated average flipping probability profiles of TF core motifs with the sequence classification for DNA binding (1 for bound and 0 for unbound).

We observed a wide range of correlation and significance values for the TFs tested, with *MAF bZIP transcription factor K (MAFK)* notably displaying the highest positive correlation, and *Zinc finger and BTB domain containing 7A (ZBTB7A)* exhibiting the most negative correlation with binding affinity (Fig 4a). After Bonferroni correction (*ɑ* < 0.05), we found that 39 out of 44 TFs showed statistically significant correlations. We highlighted the results from *MAFK* and *ZBTB7A* here. In *MAFK*-bound sequences, the average flipping probability in TF motifs is substantially higher than in unbound ones (Fig 4b). Conversely, for *ZBTB7A*, unbound sequences had higher average flipping probability. Together these results suggest that for a substantial proportion of commonly studied TFs, their binding to DNA may be influenced by DNA breathing, however, likely using different structural mechanisms.

**Fig. 4:**
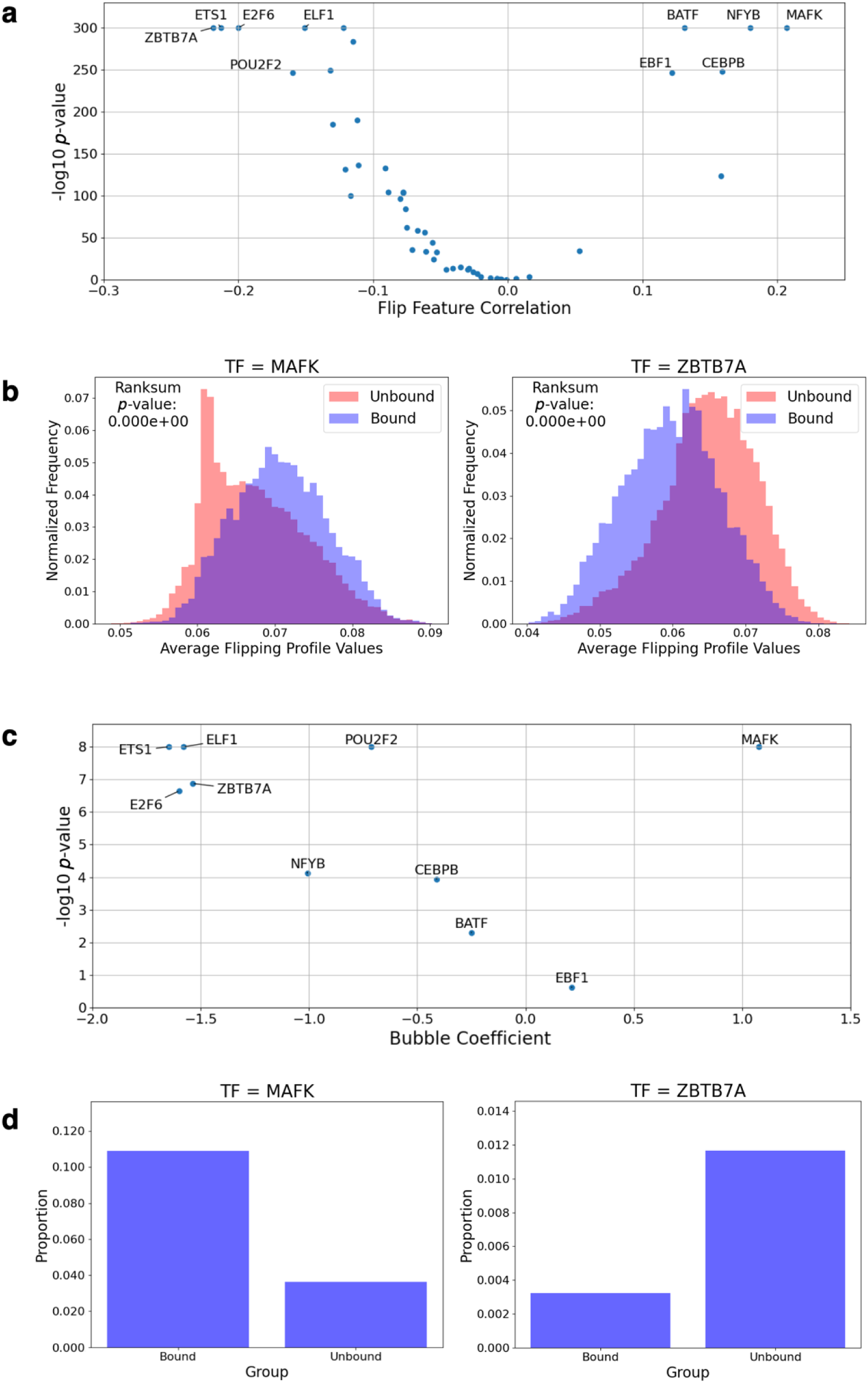
Analysis results of tests performed on a ChIP-seq dataset using 40 TFs. (a) Correlation coefficients for average flipping probability profile values at motif positions plotted against the correlation significance. (b) Histogram for the average flipping probability profile values at motifs in a sequence split according to the TF being bound (binding status = 1) or unbound (binding status = 0), with the highest positive correlation observed for *MAFK* and highest negative correlation observed for *ZBTB7A*. (c) Correlation coefficients derived through logistic regression for the distance between a motif and the nearest bubble, with distance defined as 1 if distance ≤ 10 bp, and 0 if distance > 10 bp, plotted against the correlation significance. (d) Proportion of sequences with a bubble ≤ 10 bp away from a motif in bound and unbound sequences for *MAFK* and *ZBTB7A*.

We next explored the role of DNA breathing using DNA bubble propensity. Computation of bubble propensity features, however, relies on compute-intensive molecular simulations, so we limited this analysis to the 10 TFs from the flipping probability analysis above, including the top five positively correlated TFs and top five negative control ones. For each sequence, we select the strongest motif match (based on *p*- value). We then tested if the presence of a bubble nearby (within 10 bp of the motif) correlated with TF binding status (1 or 0). For all five TFs whose flipping probability showed negative correlations with TF binding, their bubble propensity feature also correlated negatively with TF binding (Fig 4c). For the five TFs whose flipping probability showed positive correlations, with varying patterns (Fig 4c). *MAFK* showed a strongly positive correlation. *Nuclear transcription Factor Y subunit Beta (NFYB)* and *CCAAT/Enhancer-Binding Protein Beta (CEBPB)* showed modestly negative correlations while *Basic leucine zipper ATF-like transcription factor (BATF)* and *Early B-cell Factor 1 (EBF1)* showed insignificant correlations. We also repeated the analysis with different distance cutoffs to define nearby bubbles (5 bp, 20 bp), and the results were essentially similar (S3 Fig).

Examining *MAFK* and *ZBTB7A* in more detail, we found that for *MAFK*, a higher proportion of bound sequences contain bubbles near motifs (about 3-fold enriched, Fig 4d). In contrast, for *ZBTB7A*, unbound sequences were more likely to have bubbles close to motifs, suggesting that bubbles might interfere with binding for this TF (Fig 4d), again pointing to likely different structural readout modes at play.

Altogether, these results broadly support the relevance of DNA breathing in TF-DNA interactions. It also underscores the complexity of the role of DNA breathing in TF binding. The effects of base flipping varied significantly among different TFs, with some TFs likely benefiting from increased flipping and others being hindered. The results from bubble feature analysis generally agreed with those overserved in the flipping analysis. The disagreement of the results from some TFs could be due to the lower power of the bubble analysis, given that DNA bubbles are relatively rare. Additionally, the proximity of bubbles to motifs may play a role.

## Discussion

We presented a comprehensive study of the role of DNA breathing in TF-DNA interactions. Given that TF-DNA interactions underlie much of gene regulation, we think this work is an important advance of our current understanding of gene regulation.

Using both in vitro and in vivo TF binding data, our analyses showed that DNA flipping probability in motif positions correlated with TF binding affinities, and that DNA bubble propensity near motifs also correlated with binding. The sign and magnitude of these associations, however, seem to vary across TFs, suggesting that the role of DNA breathing may depend on specific structural feature readout mechanisms employed by various of TFs.

Our results, while being generally supportive of the role of DNA breathing, have some limitations. We found that the results using flipping probability and bubble propensity features generally agree. This is somewhat expected because a base in a single- stranded region of DNA will more likely flip. This is due to energetic considerations. A base without intra-bp hydrogen bonds with the second strand engages likely in weaker stacking interactions with its adjacent bases and can therefore flip. This said, the two features also capture somewhat different aspects of DNA breathing, with flipping probability profiles directly representing an aspect conformational flexibility and strength of inter-bp stacking whereas bubbles represent the tendency of single-strand DNA formation and thereby the simultaneous opening of intra-bp hydrogen bonds in multiple adjacent bp.

For some TFs, it is not conclusive yet whether higher or lower DNA breathing correlates with higher TF binding. Another limitation is that our results are based on the statistical pattern of DNA breathing probability. We do not yet have proof that DNA breathing causally affects TF binding. An interesting future analysis would be to use genetic variation data to show that genetic variants affecting DNA breathing (beyond motif recognition), also affect TF binding.

It is worth pointing out other directions for future studies. First, it would be interesting to develop a quantitative model of TF-DNA interactions incorporating DNA breathing. This possibility has been shown in a recent paper that used deep neural networks incorporating both DNA sequences and DNA breathing features [16]. However, this study has several limitations, as discussed in the Introduction. We caution that DNA breathing itself depends on the underlying DNA sequences, and this type of model insufficiently considered the dependence of DNA breathing on nucleotide sequence. Deep learning models are also notoriously difficult to interpret. It would be interesting to apply strategies for interpretability in protein-DNA binding [21] or develop models based on biophysical principles, for example, a model where the free energy of TF-DNA binding at any position of the motif depends on DNA breathing features at that position. It would also be interesting to explore DNA breathing from the perspective of TF families and structure. For example, using the structure of a DNA binding interface with a TF [22] may allow one to assess to what extent binding may be influenced by DNA bubbles.

These complementary approaches may help explain our finding that the role of DNA breathing seems to vary across TFs.

In conclusion, our study provided support to the role of DNA breathing properties in TF-DNA interactions and, thereby, outlines several interesting new directions for future research.

## Methods

### TF binding data from gcPBM and ChIP-seq

We investigated transcription factor (TF) binding affinities using two datasets. The first dataset, derived with gcPBM binding arrays, included 23,028 sequences, each 36 bp long, with binding affinities assigned on a scale from 0 to 1 for the bHLH TF complexes: *Max-Max (MAX)*, *Mad2-Max (MAD)*, and *c-Myc-Max (MYC)* [18]. The second dataset comprised 886,625 sequences, each 200 base pairs long, annotated with ChIP-seq peak data from the DeepSEA study [2] for 64 different TFs. Here, binding affinity was dichotomous, recorded as either 1 (peak present, bound) or 0 (peak absent, unbound).

### Computational motif scanning

We utilized FIMO from the MEME Suite [23] to perform the motif scans, using position weight matrices (PWMs) obtained from the JASPAR2022 database [24]. We categorized motif matches into two strength levels: strong (*p*-value < 1x10e-4) and weak (*p*-value between 1x10e-3 and 1x10e-4). This process allowed us to pinpoint exact bp positions containing a motif for a given TF, and a corresponding *p*-value for that motif match. Since JASPAR does not have a motif PWM for the TF MAD, it was excluded from our gcPBM dataset analysis. Similarly, 20 of the 64 TFs present in our ChIP-seq database did not have motif PWMs in JASPAR and were excluded from analysis.

### Defining DNA breathing features

Our breathing data was generated using <u>pyDNA-EPBD</u>, an MCMC simulation package for Python which uses the Extended Peyrard-Bishop-Dauxois (EPBD) nonlinear DNA model to describe DNA dynamic features [17]. The simulation settings were set to the standard recommended by the package, with the temperature set to 310 K, preheating equilibration steps set to 50,000, and simulation production steps set to 80,000. The package outputs a three-dimensional tensor of DNA bubble probabilities, where each value in the tensor is defined as the probability of a certain bubble of length *l* (measured horizontally across the DNA molecule in bp length) with displacement *thr* (measured in Å) forming at a bp position *n*. Here displacement means the distance of a base position in an open bubble from what its position would be if the bubble were closed.

The flipping probability feature is defined as probability that a specific bp is flipped out of the double helix, and a given bp is considered flipped based on a certain displacement threshold in Å. We calculated flipping probability at a displacement threshold of 0.5 * √2 Å. Due to the sheer volume of ChIP-seq sequences used to compute the flipping probability correlations in our analysis, we used features generated by an external collaborator. For the gcPBM sequences, we computed flipping probability ourselves since it was computationally manageable. The simulation also generates coordinate distance values (as well as the square of that value) for each bp in the sequence given to it, which we used as alternative features to define flipping.

To define DNA bubbles, we used a threshold for bubble size, based on a paper published by the authors of the simulation package [14]. A bubble has displacement > 3.5 Å and bubble length > 10 bp. Using these criteria on our dataset, we were able to slice the tensor to yield one bubble probability value per bp position. We chose the 95th percentile probability value as a significant threshold to categorize whether a bubble was present at a bp position or not. This criterion resulted in ∼20% bubble-positive sequences in the gcPBM data and ∼40% in the ChIP-seq data.

### Testing correlation of flipping features with binding affinity in gcPBM data

For the first part of the correlation analysis, we averaged flipping probability across all motif positions in a sequence, giving us a single average flipping probability per sequence that we then correlated against the binding affinity of the sequences. For the second part of the analysis, we took the flipping probability at each individual bp location in a motif and correlated these values against binding affinity of the sequences to yield correlation results at each bp position of the motif.

### Testing correlation of bubble presence with binding affinity in gcPBM data

We compared binding affinity of sequences with the bubble propensity feature, which indicates the presence of a bubble (bubble positive) versus sequences without bubbles (bubble negative). We first conducted a Wilcoxon Rank Sum test for each TF to determine if the binding affinity significantly differed between the two groups. To account for the effects of motifs, we ran two sets of linear regression analysis of bubble presence against binding affinity. In the first analysis, we controlled for the presence of motifs by keeping motif presence as a covariate in the regression. In the second analysis, we limited it to sequences without strong motif matches.

### Testing correlation of flipping features with binding affinity in ChIP-seq data

We replicated the two methodologies used for the gcPBM data using 44 different TFs, with some TFs having more than one motif (PWM) with different consensus sequences.

### Testing correlation of bubbles with binding affinity in ChIP-seq data

For each sequence, we first isolated the strongest motif match (by *p*-value) in that sequence, and then assessed if there is a nearby bubble at different thresholds (5, 10, or 20 bp). We then performed logistic regressions to test the effect of bubble presence near a motif (binary variable) of a sequence on its binding affinity.

## Acknowledgements

This work was supported by the National Institutes of Health under the grants, R01MH116281 (X.H, B.A., J.D.), R01MH110531 (X.H.), and R35GM130376 (R.R). The LANL authors gratefully acknowledge the support of Los Alamos National Laboratory which is operated by Triad National Security, LLC, for the National Nuclear Security Administration of U.S. Department of Energy, Contract No.89233218CNA000001.

## Supporting Information

**S1 Table. gcPBM dataset summary table.** Summary table for number of sequences, number of sequences with a bubble present, and number of sequences with motif matches (total, strong, or weak) for the TFs in the gcPBM dataset. *MAD* does not have a motif PWM available in JASPAR and so does not have any calculated motif matches.

**S2 Table. ChIP-seq dataset summary table.** Summary table for ChIP-seq data used in the analysis of flipping probability profiles. Shown are the number of ChIP-seq labels (positive or negative for bound or unbound) and number of sequences with motif matches (total, strong, weak, strong, and weak) for the TFs in the ChIP-seq dataset (Total sequences = 886,625). TFs without a motif PWM in JASPAR do not have any calculated motif matches.

**S3 Table. Bubble propensity analysis ChIP-seq dataset subset summary table.** Summary table of the ChIP-seq data used in the bubble propensity analysis. Shown are the number of ChIP-seq labels (positive or negative for bound or unbound), number of sequences with a bubble present, and number of sequences with motif matches (total, strong, weak, strong, and weak) for the TFs in the subset of the total ChIP-seq dataset that we rendered bubble propensity features for (Total sequences = 35,331).

**S1 Fig. Defining DNA breathing features (Bubbles and Flipping).** Definition of DNA breathing feature based on simulation output. (a) DNA bubble probability values (*z*-axis) are expressed for selected amplitude thresholds in Å (figure titles), bubble propensity in number of bp (*y*-axis), and the bp position of the bubble (*x*-axis). (b) Probability values at 3.5 Å and a bubble propensity of 10 bp (highlighted in red) for a selected sequence as an example. (c) Bubble probabilities along the bp positions in the example at panel (b). Red horizontal line shows the 95% quantile of bubble probability. (d) The bp-positions with bubble probability values above the significant threshold, in panel (c), were considered to indicate that a bubble is present at that position, allowing us to define bubble presence and location in a sequence. (e) DNA flipping probability values were generated at five distance thresholds, with each threshold being a multiple of 0.5 * √2 Å. (f) We selected the flipping probability generated at the smallest threshold (highlighted in red). (g) These flipping probability values were used downstream.

**S2 Fig. Correlations of DNA bp coordinate displacement values and the square of those values at motif locations with respective binding affinity.** Correlations of DNA bp coordinate displacement values and the square of those values at motif locations with respective binding affinity. (a) The average coordinate displacement values of motif positions in a sequence were correlated against binding affinity values of sequences for MAX and MYC. (b) The average coordinate displacement values at each position in the motif for MAX and MYC were independently correlated with binding affinity of the sequence. Bonferroni corrected correlation significance with *ɑ* = 0.5 is denoted by a red asterisk. (cd) The same analysis as reported in (ab), using *r^2^* of the coordinate displacement values.

**S3 Fig. Correlation of bubble presence nearby a motif with binding affinity.** Correlation of bubble presence nearby a motif with binding affinity. Results using distance cutoffs of 5 bp (panel a) and 20 bp (panel b), respectively.

